# Transcriptomic response during wasp parasitism in the *Drosophila*-*Spiroplasma* interaction

**DOI:** 10.1101/2020.08.27.270462

**Authors:** Victor M. Higareda-Alvear, Mariana Mateos, Diego Cortez Quezada, Cecilia Tamborindeguy, Esperanza Martínez-Romero

**Affiliations:** Center for Genomic Sciences, Universidad Nacional Autónoma de México, Cuernavaca, Morelos, México; Department of Ecology and Conservation Biology, Texas A&M University, College Station, TX, USA; Department of Entomology, Texas A&M University, College Station, TX, USA

**Author notes:** Design and conceptualization: VMHA, MM, CT Conducted experiments: VMHA, MM Resource acquisition: MM, CT, EMR Performed analyses: VMHA, MM Interpreted Results: VMHA, MM, DCQ, CT, EMR Prepared figures and tables: VMHA Wrote first draft: VMHA, MM Revised, edited, approved manuscript: VMHA, MM, DCQ, CT, EMR.

**Keywords:** Metatranscriptome, Spiroplasma, Parasitic wasp, Protection, Immunity, Drosophila, Toxins

## Abstract

Several facultative bacterial symbionts of insects protect their hosts against natural enemies. *Spiroplasma poulsonii* strain *s*Mel, a male-killing heritable symbiont of *Drosophila melanogaster*, confers protection against some species of parasitic wasps. Several lines of evidence suggest that *Spiroplasma*-encoded ribosome inactivating proteins (RIPs) are involved in the protection mechanism, but the potential contribution of the fly-encoded functions has not been deeply explored. Here we used RNA-seq to evaluate the response of *D. melanogaster* to infection by *Spiroplasma* and parasitism by the *Spiroplasma*-susceptible wasp *Leptopilina heterotoma*, and the *Spiroplasma*-resistant wasp *Ganaspis hookeri*. In the absence of *Spiroplasma* infection, we found evidence of *Drosophila* immune activation by *G. hookeri*, but not by *L. heterotoma*, which in turn negatively influenced functions associated with male gonad development. As expected for a symbiont that kills males, we detected extensive downregulation in the *Spiroplasma*-infected treatments of genes known to have male-biased expression. We detected very few genes whose expression was influenced by the *Spiroplasma-L. heterotoma* interaction, and they do not appear to be related to immune response. For most of them, parasitism by *L. heterotoma* (in the absence of *Spiroplasma*) caused an expression change that was at least partly reversed when *Spiroplasma* was also present. It is unclear whether such genes are involved in the *Spiroplasma*-mediated mechanism that leads to wasp death or fly rescue. Nonetheless, the expression pattern of some of these genes, which reportedly undergo expression shifts during the larva-to-pupa transition, is suggestive of an influence of *Spiroplasma* on the development time of *L. heterotoma*-parasitized flies. In addition, we used the RNAseq data and quantitative (q)PCR to evaluate the transcript levels of the *Spiroplasma*-encoded RIP genes. One of the five RIP genes (RIP2) was consistently highly expressed independently of wasp parasitism, in two substrains of *s*Mel. Finally, the RNAseq data revealed evidence consistent with RIP-induced damage in the ribosomal (r)RNA of the *Spiroplasma*-susceptible, but not the *Spiroplasma*-resistant, wasp. We conclude that immune priming is unlikely to contribute to the *Spiroplasma*-mediated protection against wasps, and that the mechanism by which *G. hookeri* resists/tolerates *Spiroplasma* does not involve inhibition of RIP transcription.

## 1 Introduction

During their life cycle, insects face a large diversity of natural enemies such as predators and parasites, as well as infections by bacteria, fungi, and viruses. Although insects rely on an immune system to overcome these infections (reviewed in Hillyer, 2016), parasites and pathogens have evolved counter defenses. In this arms race, many insects have allied with symbiotic bacteria to fight against parasites. Extensive evidence of such defensive symbioses has been accrued over the last ~17 years (Oliver and Perlman, 2020).

Three models of classical ecology can be adapted to explain protection of bacteria against parasites. **Exploitation competition** occurs when the symbiont and the parasite compete for a limiting resource (e.g. *Wolbachia* and vectored-viruses compete for cholesterol;(Caragata et al., 2013). **Apparent competition** can occur when the symbiont activates (“primes”) the immunity of the host, and thus indirectly interferes with the parasite (e.g. *Wigglesworthia* in *Glossina* against trypanosomes, (Wang et al., 2009). Finally, **interference competition** can occur when the symbiont produces a compound (e.g. a toxin) that limits the success of the parasite (e.g. *Hamiltonella defensa* in aphid insects (Oliver et al., 2003; Brandt et al., 2017). One or more of these mechanisms can occur in concert, as suggested for the interactions between flies in the genus *Drosophila* and heritable bacteria in the genus *Spiroplasma*, where these endosymbionts protect the host against parasitic wasps or nematodes (Jaenike et al., 2010; Xie et al., 2010).

The association between *Drosophila melanogaster* and its naturally occurring heritable bacterium *Spiroplasma poulsonii* (*s*Mel) has emerged as a model system to study the evolutionary ecology and mechanistic bases of both defensive mutualisms (Xie et al., 2010; Ballinger and Perlman, 2017) and reproductive parasitism (i.e, male-killing) (Cheng et al., 2016; Harumoto et al., 2016).

The presence of *Spiroplasma* in *Drosophila* larvae prevents the successful development of several species of parasitic wasps (hereafter “*Spiroplasma*-susceptible” wasps) that oviposit on larvae, which leads to enhanced fly survival in some interactions, but not others. Furthermore, several wasp species are unaffected by the presence of *Spiroplasma* in the host, which we hereafter refer to as “resistant” (Mateos et al., 2016). The degree of *Spiroplasma*-mediated protection varies by wasp genotype (Jones and Hurst, 2020b) and is dependent on abiotic factors, e.g. temperature (Corbin et al., 2020)

Research into *Spiroplasma*-mediated protection against wasps has revealed that *Spiroplasma*-susceptible wasp embryos manage to hatch into first instars and achieve some growth, which is subsequently stalled (Xie et al., 2010, 2014; Paredes et al., 2016). Evidence consistent with competition for lipids (i.e., exploitative competition) between *Spiroplasma* and the developing wasp has been reported for the wasp *Leptopilina boulardi* (Paredes et al., 2016) Regarding the role of interference competition, the genomes of several *Spiroplasma* strains, including *s*Mel, encode genes with homology to ribosome inactivating proteins (RIP’s, Ballinger and Perlman, 2017).

RIP proteins, which are produced by different plants and bacteria (e.g. ricin and Shiga toxin, respectively), cleave a specific adenine present within a highly conserved (i.e., in all eukaryotes) motif of the large ribosomal subunit (28S rRNA), leading to inactivation of the ribosome and inhibition of protein translation (reviewed in Stirpe, 2004).

Damage consistent with RIP activity (hereafter referred to as depurination) has been detected in a *Spiroplasma*-susceptible nematode (Hamilton et al., 2015) and in two *Spiroplasma*-susceptible wasps (the larval parasitoids *L. boulardi* and *L. heterotoma*), but not in a *Spiroplasma*-resistant wasp that oviposits on fly pupae (Ballinger and Perlman, 2017).

Although the above studies suggest that competition for nutrients and RIP activity are involved in the *Spiroplasma*-mediated mechanism that causes wasp death, they have not demonstrated that the above mechanisms alone or in combination are necessary and sufficient, and alternative mechanisms, including immune priming, have not been ruled out.

In response to wasp parasitism, *D. melanogaster* mount an inmune response characterized by proliferation of blood cells also known as hemocytes. Plasmatocytes are the first cells to attach to the foregin egg followed by lamellocytes which form successive layers; both types of hemocytes consolidate around the wasp egg, forming a capsule. The inner cells of the capsule produce melanin and release free radicals into the capsule, killing the wasp (Russo, 1996; Carton et al., 2008). Wasps have evolved a diverse array of strategies that counter the fly-encoded defense (Schlenke et al., 2007; Mortimer et al., 2013). Whether or not *Spiroplasma* contributes to enhancing the fly-encoded defense against wasps has not been extensively investigated. To date only one study has examined the possible influence of *Spiroplasma* on fly-encoded immunity against wasps (Paredes et al., 2016). Their results revealed no effect of *Spiroplasma* on the number of hemocytes in flies parasitized by *L. boulardi*. Whether *Spiroplasma* influences this or other aspects of fly-encoded immunity against other wasps has not been examined.

Herein, we used an RNA-seq approach to evaluate the transcriptomic response of *D. melanogaster* during interactions involving *Spiroplasma* and two wasps that are generalists of the genus *Drosophila*: the *Spiroplasma*-susceptible *L. heterotoma*; and the *Spiroplasma*-resistant *Ganaspis hookeri*. In addition, we evaluated the effect of wasp parasitism on the expression of *Spiroplasma* RIP genes in two closely related substrains of *s*Mel, which have similar genomes (Gerth et al., 2020) but confer different levels of protection against wasps, measured as fecundity of flies surviving a wasp attack (Jones and Hurst, 2020a).

## 2. Methods

### 2.1 Insect and *S. poulsonii* strain

The transcriptomic experiments were performed on *D. melanogaster* flies (strain Canton S), which naturally harbor *Wolbachia* (Riegler et al., 2005). Flies were reared in a Standard cornmeal medium (recipes in Supplementary methods 1) at 25°C, with a dark: light 12 hour-cycle. Canton S flies were artificially infected with *Spiroplasma poulsonii* strain *s*Mel-BR (original isofemale line “Red42” from Brazil; (Montenegro et al., 2005) via hemolymph transfer (as in Xie et al., 2010) at least 3 generations before initiating the experiment. As *s*Mel-BR is a male-killer, the *Spiroplasma-*infected strain was maintained by addition of *Spiroplasma*-free males (Canton S strain) every generation. Wasps, *Leptopilina heterotoma* strain Lh14 (Schlenke et al., 2007) and the all-female *Ganaspis hookeri* strain G1FL (Mortimer et al., 2013), were reared using second instar Canton S *Spiroplasma*-free larvae. These wasp strains are naturally infected with one or more *Wolbachia* strains (Wey et al., 2020 and Mateos, unpublished data). For the qPCR assays, we used *Wolbachia*-free Oregon R flies to which *s*Mel-BR or *s*Mel-UG ( original isofemale line from Uganda, Pool et al., 2006), had been artificially transferred at least 3 generations prior. These flies were maintained by matings with *Spiroplasma*-free Oregon R males under the same environmental conditions as the Canton S background flies, but in a opuntia-banana food medium (recipes in Supplementary methods 1).

### 2.2 RNA-Seq based methods

#### 2.2.1 Wasp exposure

To examine the effect of the interaction of *Spiroplasma* and wasp on the transcriptome of *Drosophila* (and of *Spiroplasma*), we compared treatments with the presence and absence of *Spiroplasma* (*s*Mel-BR) and one of the two wasp species at two different time points. This experimental design resulted in a combination of twelve treatments; six treatments per time point. For each replicate, parental flies (approximately ten females and ten males) were set up in oviposition vials in the evening for overnight oviposition. Parental flies were removed the next morning. One day later, ~30 second-instar *D. melanogaster* larvae were carefully collected and transferred to a Petri dish (60 mm diameter) containing cornmeal food medium (**Supplementary methods 1**). In replicates assigned to a wasp treatment, 5 male and 6 female wasps of the corresponding species (*L. heterotoma* or *G. hookeri*) were added to the Petri dish, and allowed to oviposit for ~5 h. All female wasps had been previously allowed to oviposit on *D. melanogaster* fly larvae for ~5 hours. The purpose of this “training” is to ensure that the wasps are experienced at oviposition prior to the experiment. All Petri dishes were covered, but a small hole was opened (with a hot needle) to allow for gas exchange, and/or through which wasps could feed on a piece of cotton wool soaked in 1:1 water:honey mix that was placed outside the dish. To collect RNA, larvae were retrieved from each Petri dish at either 24h (T1) or 72h (T2) post-wasp attack (PWA) (i.e., one or three days after wasps were removed, respectively). To ensure sufficient material for RNA-seq, 20–30 larvae were collected for the 24h time-point, whereas 10–20 for the 72h time point. Both wasp species were embryos at T1 and larvae at T2. Fly larvae from the same Petri dish were pooled into a single RNA extraction tube (i.e., a replicate).

From each Petri dish in the wasp-exposed treatments, a subsample of fly larvae was used to verify wasp parasitism rate as follows. First, at the time of larvae collection for RNA, five fly larvae per replicate were dissected under the microscope and discarded. If all five larvae contained at least one wasp egg or larva (i.e., 100% wasp oviposition rate), the replicate was retained and processed. If one or more fly larvae did not contain a wasp larva (or embryo), then five additional fly larvae were examined for wasp presence. Only replicates with 90-100% wasp oviposition rate were retained. All collections for RNA were performed in the afternoon-evening, and collected larvae were quickly placed in an empty microtube for processing.

#### 2.2.2 RNA extraction

Collected larvae were either immediately frozen at −80°C or immediately processed for RNA extraction. Preliminary experiments with RNAlater Stabilization Solution revealed that the larvae did not die immediately and appeared to melanize. Total RNA was extracted using the Trizol [invitrogen] method. Each sample consisted of a pool of *Drosophila* larvae that were homogenized by hand with a sterile plastic pestle in Trizol reagent. The Trizol isolation method was performed following the manufacturer’s protocol but it was stopped at the 70% ethanol wash step. The total RNA pellet in ethanol was submitted to the Texas AgriLife Genomics and Bioinfomatics Services facility for completion of the RNA isolation procedure, assessment of RNA quality and quantity (using Fragment Analyzer; Agilent, Santa Clara, CA), library preparation, sequencing, and demultiplexing.

#### 2.2.2 Library preparation and sequencing

Total RNA was subjected to removal of ribosomal RNA from eukaryotes and prokaryotes with the RiboZero Epidemiology kit (Illumina, San Diego, CA). The TruSeq stranded kit (Illumina) was then used to prepare the library for sequencing with Illumina (125 bp Single End “HighSeq 2400v4 High Output”).

#### 2.2.4 Bioinformatic analysis

Quality and presence of adapters was evaluated with FastqC (Andrews S., 2010), followed by a trimming with Trimmomatic v.0.36 (Bolger et al., 2014) using ILLUMINACLIP:/adapters.fasta:2:30:10 LEADING:3 TRAILING:3 SLIDINGWINDOW:4:15 MINLEN:36. To examine differential expression of *D. melanogaster* genes, trimmed reads were mapped with Hisat2 v.2.0.2-beta (Kim et al., 2015), using --rna-strandness R option. Treatments parasitised by *L. heterotoma* were mapped to an index composed by *D. melanogaster* genome (ensembl version BDGP6) plus *L. heterotoma* genome, reference VOOK00000000. Treatments parasitised by *G. hookeri* were only mapped to *Drosophila*, because there is not an available genome for *G. hookeri*. The resulting *D. melanogaster* mapped reads were quantified using featureCounts from the Subread package v1.6.2 (Liao et al., 2014), using the following parameters: -s 2, -t exon, -g gene_id. Differential gene expression was done in R (R core Team) with edgeR v 3.24.3 package (Robinson et al., 2009). Genes with counts < 1 cpm for all replicates and treatments under comparison were discarded. Only genes with absolute 2LogFC >= 0.58 and FDR < 0.05 were considered differentially expressed (DE). The Robust parameter robust=TRUE) of edgeR was implemented to minimize false positive DE genes. The gene ids corresponding to the DE genes were loaded (before May 2020) into Flymine, (an integrated database for Drosophila genomics(Lyne et al., 2007) available on the website https://www.flymine.org/flymine/begin.do. This platform outputs gene names plus other information such as Gene Ontology (GO), enrichment of pathways, tissue expression and protein domains. Expression patterns of individual genes were obtained from flybase2.0 (Thurmond et al., 2019) available flybase.org/. Venn diagrams used to detect exclusive genes in the interactions were generated using one web-tool available at http://bioinformatics.psb.ugent.be/webtools/Venn/

To measure *Spiroplasma* gene expression, the trimmed reads from the *Spiroplasma*-infected treatments were subjected to kallisto v.0.43.1 (Bray et al., 2016), using the genome of *S. poulsonii s*Mel-UG as reference GCF_000820525.2). Count tables obtained by kallisto were used to detect DE genes using edgeR. Expression of RIP in transcriptome was obtained from this table using trimmed mean of M-values (TMM) normalized counts. Heatmaps of expression patterns were generated with the R package pheatmap v 1.0.12.

#### 2.2.5 Power analyses for differential expression

To identify potential limitations on the detection of DE genes, we performed a statistical power analysis with the R package RNASeqPower v.1.22.1 (Hart et al., 2013). Parameters used to run different simulations were inferred from our datasets and are provided in Table S1 and Figure S1.

#### 2.2.6 Analyses of depurination signal in the sarcin-ricin loop (SRL) of the 28S rRNA of wasps and flies

RIP toxins remove a specific adenine present in the sarcin-ricin loop (SRL) of the 28S rRNA leaving an abasic site (i.e., the backbone remains intact). When a reverse transcriptase encounters an abasic site, it preferentially adds an adenine in the nascent complementary DNA strand. This property, which results in an incorrect base at the RIP-depurinated site in the cDNA and all subsequent PCR amplification steps, has been used to detect evidence of RIP activity in any procedure that relies on reverse transcription such as RNA-seq or reverse-transcription qPCR (e.g. Hamilton et al., 2015).

To examine whether a signal of depurination consistent with RIP activity was detectable in wasp-derived sequences, we mapped RNA-seq data to a reference sequence file comprised of the 28S rRNA sequences of the wasps and of *D. melanogaster* using Bowtie2 v.2.1.054 with default options. Only sequences that mapped to the wasp 28S rRNA were retained (Dataset S1). To visualize and count the shift from A to T (or other bases), the retained reads were mapped again to the 28S rRNA of wasp in Geneious v.11.1.2 (Biomatters Inc., Newark, NJ; “low sensitivity mode”; maximum gap size = 3; iterate up to 25 times, maximum mistmatches per read 2%). The number of reads containing each of the four bases or a gap at the target site was counted by selecting the position at all the reads to be counted, and recording the counts reported by Geneious under the “Nucleotide Statistics” option (gapped reads were excluded from counts). Reads were counted only if they fully covered a specific part of the 28S loop sequence (TACG**A**GAGGAACC). The bold-faced adenine represents the site of RIP depurination. Replicates with fewer than 10 mapped reads were discarded, Table S3 and Figure S10. Statistical analysis was conducted in R v 4.02 (R core Team) using a Bayesian generalized linear model (bayesglm function in “arm” package), this is because of the presence of zeros (no depurination) in some treatments. Using the above strategy, raw sequences were also mapped to the full sequence of the 28S rRNA of D. melanogaster, and depurination was evaluated. Due to the high number of ribosomal sequences that align to the 28S rRNA, only subsets of 1 million of sequences were analyzed (Table S4).Using the above strategy, raw sequences were also mapped to the full sequence of the 28S rRNA of *D. melanogaster*, and depurination was evaluated. Due to the high number of ribosomal sequences that align to the 28S rRNA, only subsets of 1M of sequences were analyzed. Table S4

### 2.3 Quantitative (q)PCR-based Methods

#### 2.3.1 Expression of *Spiroplasma* Ribosomal inactivation proteins (RIPs)

To verify the RIP expression patterns inferred from the transcriptome (see Results) and to examine whether they were consistent between substrains of *s*Mel, we used qPCR on a a new set of treatments. We followed the “Wasp exposure” methodology (described above), but used *Wolbachia-*free *D. melanogaster* (OregonR) harboring the *S. poulsonii* strain *s*Mel-BR or *s*Mel-UG. Five larvae per treatment (parasitized or not by *L. heterotoma* or *G.hookeri*), were collected at 24 and 72h PWA, flash frozen in liquid nitrogen and homogenized by hand with a pestle. Total RNA was extracted with the All prep DNA/RNA mini kit (Qiagen, Germantown, MD). 1ug of total RNA was used to synthesize cDNA using superscript II reverse transcriptase (Invitrogen), following manufacturer’s procedures. cDNA was used as template for qPCRs, performed on a CFX96 detection system (Bio-Rad, Hercules, CA). The mix contained 5ul of iTaq Universal SYBR Green Supermix (Bio-Rad), 2.5ul water, 2ul cDNA, and 0.25ul of each primer (stock solution at 10uM). Primer sequences and efficiencies for RIP2, RIP3-5 and rpoB were taken from (Ballinger and Perlman, 2017). For RIP1, we designed and used the following primers Forward: 5’-AATCAGAGGGGCATTAGCTC-3’ Reverse 5’-CTTCGCTTGTGGTTCTTGAT-3’, efficiency = 0.995. Although (Ballinger and Perlman, 2017) reported a primer pair targeted at RIP1, this primer pair matches a fragment of RIP2 instead.

Relative expression was calculated using efficiency-corrected Ct values using (Ct*(Log(efficiency)/Log(2))) formula. DeltaCt was calculated as Ct-rpoB minus Ct-RIPx (Dataset S2). We used JMP Pro v.15 (SAS, Cary, NC) to fit a full factorial Generalized Regression (Normal distribution) model. The response variable was delta Ct Value. The independent variables (all fixed and categorical) were: RIP gene (“RIP”: RIP1, RIP2, RIP3-5), Wasp Treatment (No wasp, Lh and Gh), *Spiroplasma* strain Brazil, Uganda) Time Point (24h or 72h). Significant effects and interactions were explored with Tukey HSD tests with Least Square Means Estimates (Supplementary file S1).

### 2.4 Data availability

All raw reads generated in this project have been submitted to NCBI under the SRA accession numbers PRJNA577145. Count tables for *D. melanogaster* and *Spiroplasma* are in Dataset S3. Command lines used to run bioinformatic analyses are available in Supplementary methods 2. Data generated during this study are available at figshare https://doi.org/10.6084/m9.figshare.c.5104559.v3

## 3. Results

To examine the effect of *Spiroplasma* on *D. melanogaster* gene expression under wasp parasitism, we generated twelve-RNA-seq treatments (Figure1), with an average of 47 million quality single-end reads per sample, ~90% of these reads mapped to the *D. melanogaster* genome. A fraction of these reads mapped to ribosomal sequences of the host, which is an indication of incomplete ribodepletion (Table S2).

**Figure 1.**
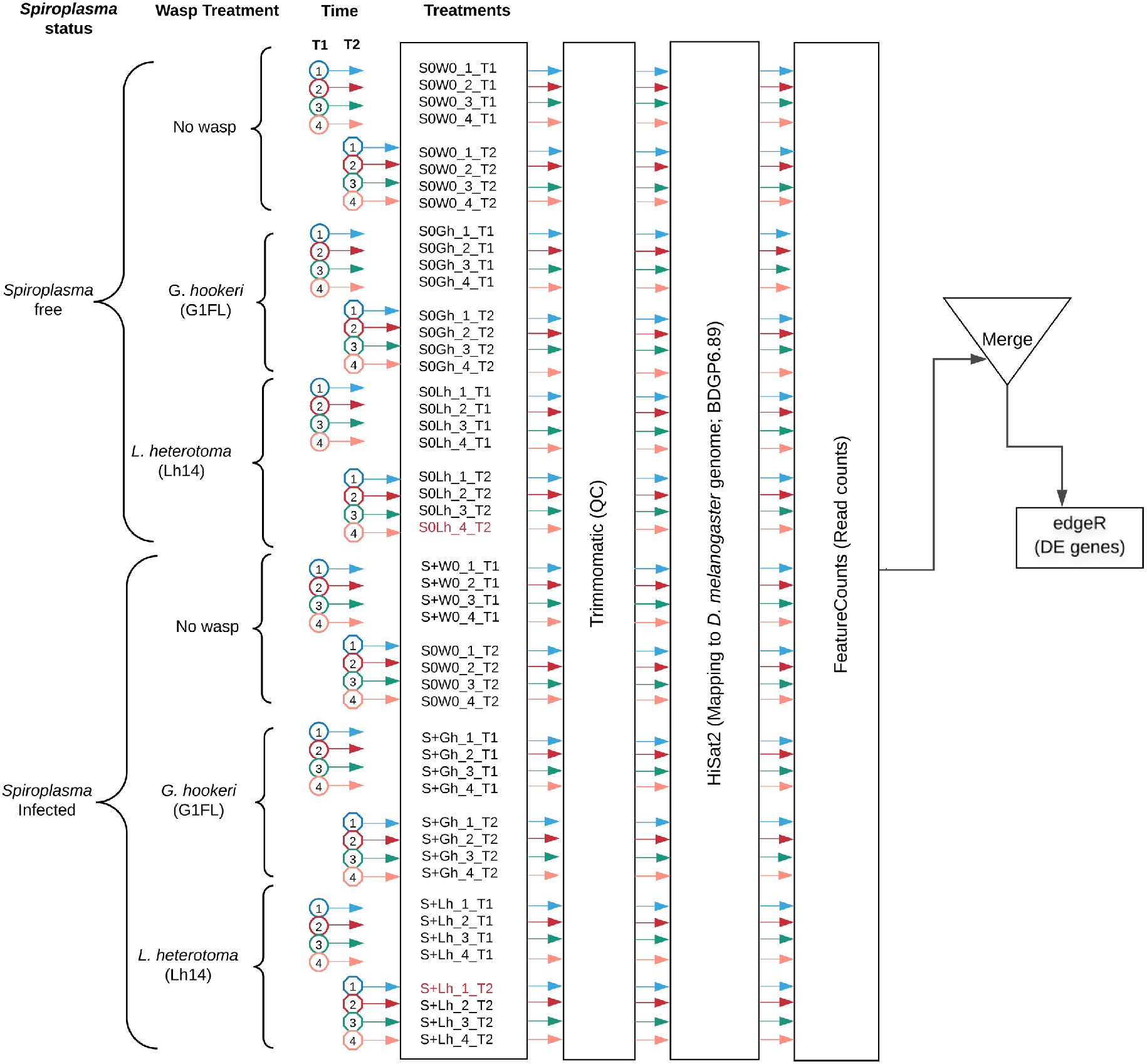
Experimental design and workflow for data analysis. Two samples (S0Lh_4_T2 and S+Lh_1_T2) were removed from the analysis (see text).

The multidimensional scaling (MDS) plot of all treatments at 24h post wasp attack (PWA; T1) did not reveal any particular grouping by treatment (Fig. S2). At the 72h PWA time point (T2), however, the treatments separated at the first dimension by presence/absence of *Spiroplasma* (Fig S3). This plot allowed us to detect two replicates that we deemed outlier and decided to exclude from further analyses (red samples in Fig 1). Replicate S0Lh_4, which lacked *Spiroplasma*, grouped with the *Spiroplasma*-infected treatments. We confirmed the absence of *Spiroplasma* because no reads derived from *Spiroplasma* were detected. Nonetheless, we realized that this replicate only contained three larvae individuals, which by chance may have been all female. Concerning replicate S+Lh_1, its expression pattern included upregulation of numerous genes associated with immune response. Consequently, we suspected that this particular replicate likely contained one or more larvae infected by a pathogenic bacterium. Below we first describe the response of *D. melanogaster* to parasitism by the wasps (*L. heterotoma* or *G. hookeri*) in the absence of *Spiroplasma*, followed by the fly response to the sole presence of *Spiroplasma*. Finally, taking these results into account, we examine the response of *Drosophila* during the *Spiroplasma*-wasp interaction.

### 3.1 Response of *D. melanogaster* to *L. heterotoma* parasitism in the absence of *Spiroplasma*

Parasitism by *L. heterotoma* at T1 (24h) post wasp attack (PWA), did not have a large effect on *D. melanogaster* gene expression, as only one gene (*thor*) was upregulated, whilst two genes, *Hml* and *mt:ATPase6*, were downregulated (Figure 2a and Dataset S5). In contrast, at T2 (72h PWA), 1216 genes were up- and 1669 down-regulated (Figure 2a and Dataset S5). Of the 1216 upregulated genes at T2, 810 grouped into 32 GO enriched categories (Figure 2b); of which transport was the most enriched. A pathway analysis revealed that upregulated genes are involved with energy generation pathways, such as lipid metabolism and citric acid cycle (Dataset S5). Of the 1669 downregulated genes, 925 grouped into 25 GO categories, of which proteolysis was the most enriched (Figure 2B and Dataset S5). Unexpectedly, a subset of 67 downregulated genes belongs to GO categories related to spermatogenesis.

**Figure 2.**
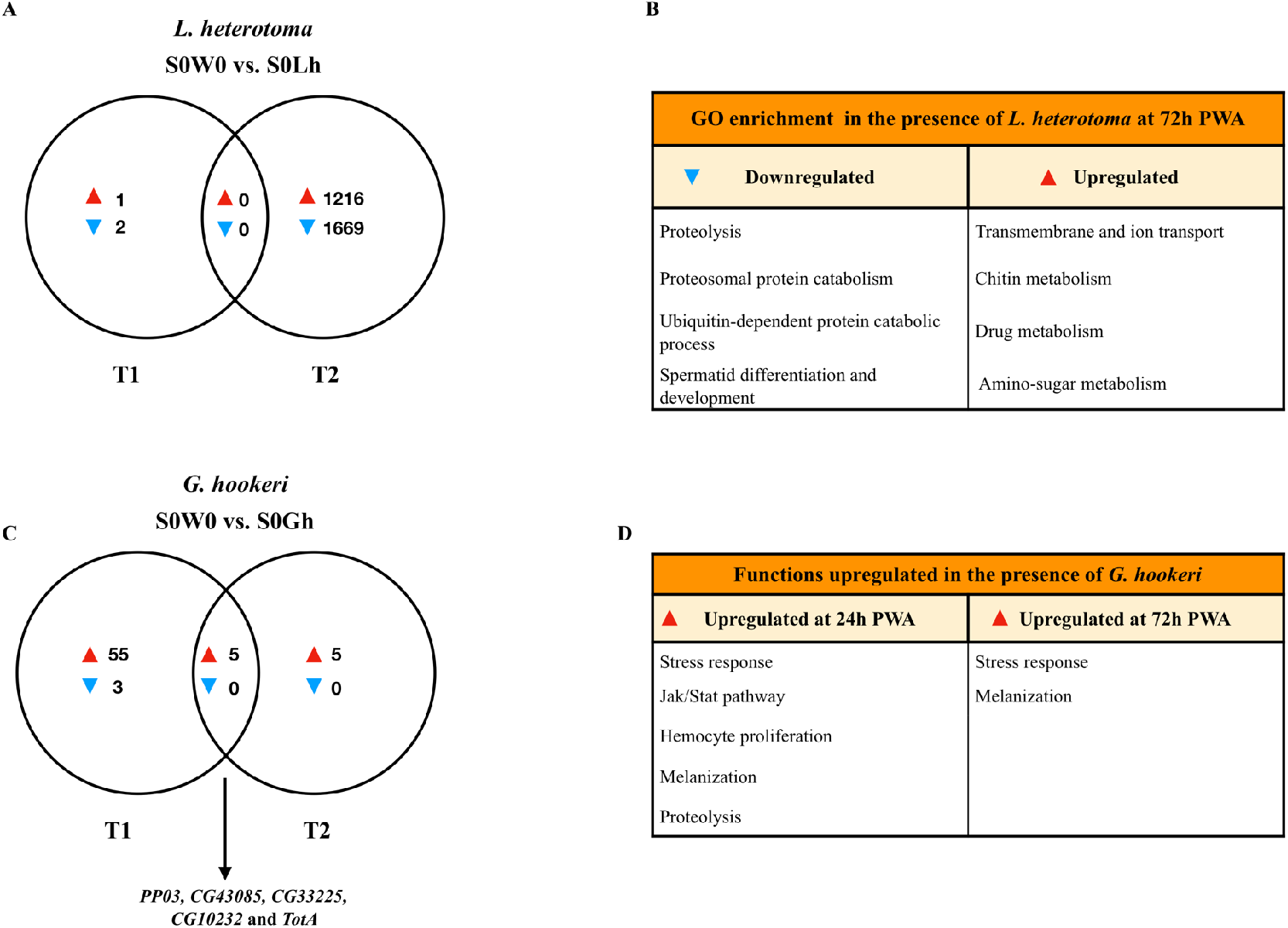
Venn diagrams and enriched functions or Gene Ontology (GO) categories. Differentially expressed genes of *D. melanogaster* at time points T1 and T2 [24h and 72h post wasp attack (PWA), respectively] by (A) and (B) *L. heterotoma* or (C) and (D) *G. hookeri*. Enriched GO categories or functions, if applicable, are reported in (B) and (D). The arrow in panel (C) indicates the five genes in the T1 and T2 intersection.

### 3.2 Response of *D. melanogaster* to *G. hookeri* parasitism in the absence of *Spiroplasma*

In the presence of the wasp *G. hookeri* (i.e, S0W0 vs. S0Gh), three genes were down- and 60 were up-regulated at T1 (Figure 2C). The two downregulated genes are known to be expressed by hemocytes (*Peroxidasin* and *hemolectin*) (Irving et al., 2005), whereas the other is a small nucleolar RNA (*Uhg4*). Only one gene, *hemolectin*, was down-regulated by both *L. heterotoma* and *G. hookeri* at T1 (Dataset S5). The sixty upregulated genes in the presence of *G. hookeri* at T1 include genes that are known to be expressed preferentially by hemocytes such as *hemese*, *ItgaPS4, ItgaPS5*, *Scavenger receptor class C* (*CG3212*), one serpin (*CG6687*), a serine protease (*CG6639*), as well as one gene involved in hemocyte proliferation (*pvf2*). (Irving et al., 2005). One activator of the Jak Stat pathway *upd3* and some effectors of this pathway, *TotA, tep1* and *tep2* (Agaisse and Perrimon, 2004), were also upregulated. Prophenoloxidase 3 (*PPO3)* and *yellow-f* genes, which are involved in the melanization process (Han et al., 2002; Dudzic et al., 2015), were also upregulated. *PPO3* was highly upregulated (2 LogFC = 9). The complete list of DE genes and GO enrichment is provided in Dataset S4.

At T2, the presence of *G. hookeri* induced upregulation of ten genes, but no genes were downregulated (Figure 2C and Dataset S5). The upregulated genes included the stress response genes *TotA*, *TotB*, *TotC* and *victoria*, but also *PPO3*; log2FC of these genes ranged 3–10. Five genes were shared between the two time points (Figure 2c). Two of them, *CG10232* and *CG33225* are predicted to be involved in proteolysis, and *CG43085* has no function or prediction assigned. The other two genes were *PPO3* and *TotA*; their log2FC were higher at T1 than at T2. In general, immune functions were upregulated by *G. hookeri* parasitism at both times (Figure 2D).

### 3.3 Response of *D. melanogaster* to *Spiroplasma*

The sole presence of *Spiroplasma* (i.e., S0W0 vs. S+W0 comparison) at 24h PWA did not reveal any upregulated genes, but 27 were downregulated. Twenty of 27 downregulated genes are reported as preferentially expressed in adult testis (Dataset S4). At 72h PWA, the presence of *Spiroplasma* induced upregulation of 16 and downregulation of 1476 genes. Only downregulated genes (692 of the 1476) were assigned to one or more of 71 GO categories (Dataset S5). Some of these categories are related to the energy generation process such as oxidative phosphorylation, pyruvate metabolic process, and glycolytic process. Among the most enriched categories were male gamete generation and spermatogenesis, which is in agreement with the expected lack of males in S+W0 treatments. In addition, 1333 of the 1476 downregulated genes at T2 are classified as preferentially upregulated in fly testis. Furthermore, the cpm values of ~78% of these genes in all replicates of the S+W0 treatment were < 1, implying very low expression levels. Therefore, as expected, a large number of the genes with lower expression in the *Spiroplasma* treatment, including the roX1 and roX2 (exclusively expressed in males, and part of the dosage compensation system; reviewed in Lucchesi and Kuroda, 2015), may simply reflect the absence of males.

### 3.4 *Drosophila* gene expression during *Spiroplasma-L. heterotoma* interaction

To explore if the presence of *Spiroplasma* influences gene expression of *D. melanogaster* during parasitism by *L. heterotoma*, we adopted the following strategy to identify genes whose expression was specifically influenced by the *Spiroplasma-L. heterotoma* interaction. We used a Venn diagram depicting differentially expressed (DE) genes from the following three pairwise treatment comparisons: S0Lh vs. S+Lh (i.e., effect of *Spiroplasma* in the presence of Lh; green sets in Fig. 3); S0W0 vs. S+W0 (i.e., effect of *Spiroplasma* in the absence of any wasp; blue sets); and S0W0 vs. S0Lh (i.e., effect of Lh in the absence of *Spiroplasma*; red sets). Genes that fell in the exclusive part of the green set were deemed as influenced specifically by the *Spiroplasma-L. heterotoma* interaction. Furthermore, genes that fell in the intersection of the green set with one or both of the blue and red sets, were further evaluated with heatmaps depicting expression levels in four treatments (S0W0, S+W0, S0Lh, S+Lh). This approach aimed to identify genes that had substantially different expression levels in the S+Lh treatment vs. the treatments that only had Lh or only had *Spiroplasma*.

**Figure 3.**
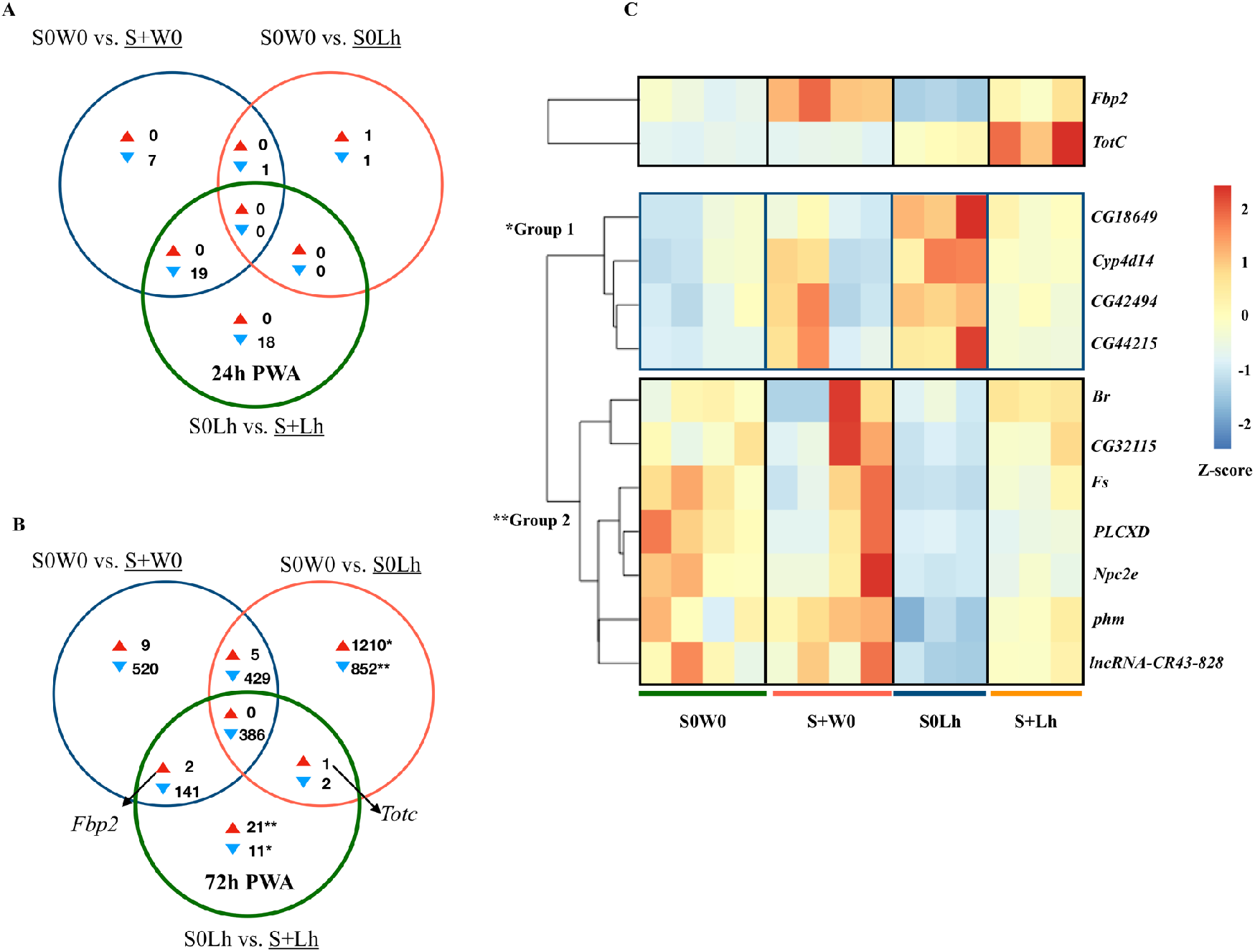
Patterns of differential expression (DE) in the context of the interaction of *Spiroplasma* and the wasp *L. heterotoma* (Lh14). Unique and shared number of DE genes in each of three treatment comparisons. (A) Time-point 1 (T1): 24h post-wasp attack (PWA). (B) Time-point 2 (T2): 72h PWA. Red- and blue-filled triangles indicate up- and downregulation, respectively, in the treatment that is underlined. (C) Heatmap of genes with peculiar expression pattern; genes in Group 1 were upregulated in the presence of only Lh and “restored” in the presence of *Spiroplasma* and Lh. Genes in Group 2 were downregulated in the presence of only Lh and “restored” in the presence of *Spiroplasma* and Lh (see text for explanation regarding TotC and Fbp2). Z-score of edgeR TMM-normalized values are represented.

The S0Lh vs. S+Lh comparison at T1 yielded zero up- and 18 down-regulated genes (exclusive green set in Fig. 3A, Dataset S4). Ten of these genes code for small nucleolar rnas (snoRNA). Five other genes (*fest*, *CG10063*, *SkpC*, *eIF4E3* and *cona*), albeit exclusively downregulated in *Spiroplasma-L. heterotoma* interaction, seem to be influenced by *Spiroplasma* alone, based on the observation that these genes were downregulated by *Spiroplasma* alone at the later time point T2 (Dataset S5). Furthermore, *fest* and *eIF4E3* are upregulated in testis, and thus their downregulation may simply reflect absence of males.

The only gene-containing intersection (green+blue), had 19 genes, of which 16 are preferentially expressed in adult testis (Dataset S4). The heatmap of all genes in the intersection (Fig. S4), revealed that most of them had similarly low levels of expression in the two *Spiroplasma*-infected treatments (i.e., S+W0 and S+Lh), compared to the S0W0, which had the highest. In turn, the S0Lh treatment exhibited an intermediate expression level (Fig. S4).

At T2, 21 genes were up- and 11 down-regulated exclusively in S+Lh treatment compared to S0Lh (exclusive green set in Figure 3B; Dataset S5). Four (*CG18649, Cyp4d14, CG42494* and *CG44215*) out of the 11 genes exclusively downregulated in the S+Lh treatment were exclusively upregulated in S0Lh with respect to S0W0 (i.e., identified with * and labeled “Group 1” in Fig. 3c). Seven (*Br*, *CG32115*, *Fs*, *PLCXD*, *Npc2e*, *phm* and l*ncRNA-CR43828*) of the 21 genes exclusively upregulated in the S+Lh treatment were exclusively down-regulated in S0Lh with respect to S0W0 (i.e., identified with ** and labeled “Group 2” in Fig. 3c). Hereafter, we refer to genes in Groups 1 and 2 as “restored” because parasitism by *L. heterotoma* (S0Lh) increased and decreased (respectively) their expression with respect to the S0W0 control, but in the presence of *Spiroplasma* plus *L. heterotoma* (i.e. the S+Lh treatment), expression levels appear to return to those observed in the control (S0W0; Fig.3C). In other words, *Spiroplasma* appears to “buffer” or counter the effects caused by the presence of *L. heterotoma*. Similarly, *Fbp2*, at the blue+green intersection, exhibits a similar expression pattern to genes in Group 2 (Fig. 3C). *TotC* (in the green+red intersection) also exhibited a peculiar expression pattern (Fig. 3C); it was upregulated by *L. heterotoma* parasitism (S0Lh), but its expression was highest in the S+Lh treatment.

The rest of the DE genes exclusive to *Spiroplasma-L. heterotoma* interaction were 14 up- and 7 down-regulated (Fig. 3B green set and Dataset S5). These genes did not group into any GO or specific pathway. Among the upregulated genes, there were *Amylase proximal*, *anachronism*, and the antimicrobial peptide *Attacin-C* (*AttC*; FC= 1.5), which was the only gene associated with immune response. Downregulated genes included two snoRNA (*CR34643* and *CR34560*), *HemK1* and a long non-coding RNA (lncRNA:CR32661) (Dataset S5).

The heatmaps of the red+blue+green (386 genes) and of the blue+green (143 genes) intersections at T2 (Fig. 3B and Fig S5-6) revealed the following general pattern that was also observed for the 19 downregulated genes in the blue+green intersection of T1 (see above): the two *Spiroplasma* treatments (S+W0 and S+Lh) had the lowest expression levels; the S0W0 treatment had the highest; and the S0Lh treatment was intermediate.

### 3.5 *Drosophila* gene expression during *Spiroplasma*-*G. hookeri* interaction

To identify fly genes specifically influenced by the *Spiroplasma-G*. *hookeri* interaction, we adopted the same Venn diagram plus heatmaps strategy as with the *Spiroplasma-L. heterotoma* interaction, except that the *L. heterotoma* treatments were replaced with *G. hookeri* treatments (see Fig. 4). Excluding genes induced by the sole presence of *Spiroplasma* or *G. hookeri*, resulted in eight and 21 exclusively up- and down-regulated (respectively) genes at T1 (exclusive green set, Fig 4A, Dataset S4). The eight upregulated genes were: four snoRNA (*CR33662*, *CR34611*, *CR34616*, and *CR34631*), one small nuclear RNA (*CR32162*), lysozyme E, diphthamide methyltransferase (*Dph5*) and mitochondrial ribosomal protein S14 (*mRpS14)*. Among the 21 downregulated genes were *DnaJ-1* (whose product is a cofactor of heat shock proteins), *starvin* (which acts as co-chaperone of Hsp70 proteins), *nervana 3* (which codes for one subunit of a sodium-potassium pump), *nanos* (which encodes a ribosomal RNA-binding protein), and one long non-coding RNA (*CR31400*).

**Figure 4.**
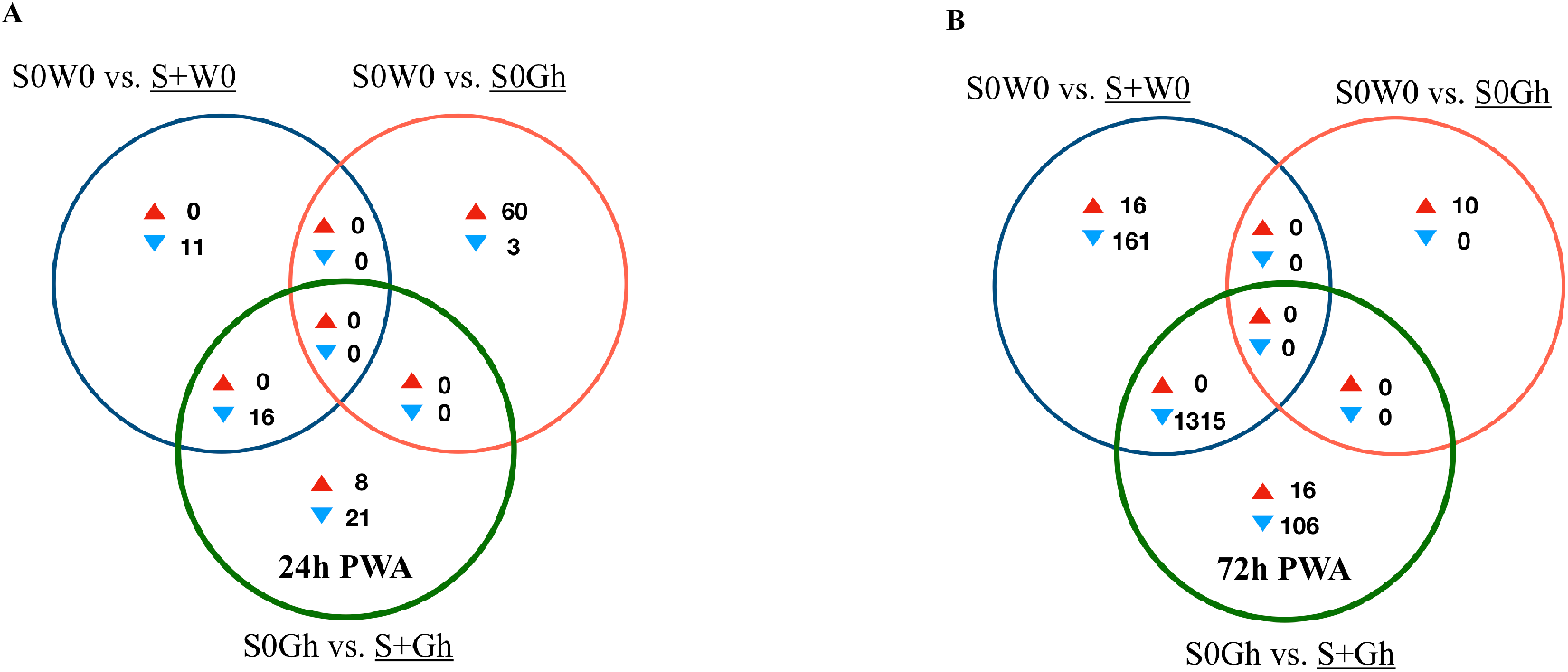
Patterns of differential expression (DE) in the context of the interaction of *Spiroplasma* and the wasp *G. hookeri* (G1FL). Unique and shared number of DE genes in each of three treatment comparisons. (A) Time-point 1 (T1): 24h post-wasp attack (PWA). (B) Time-point 2 (T2): 72h PWA. Red and blue filled triangles indicate up- and downregulation, respectively, in the treatment that is underlined.

At T1, only one intersection (green+blue) contained genes; all downregulated (n = 16; Fig. 4A). A heatmap of expression levels of these 16 genes (Fig. S7) reveals that the two *Spiroplasma* treatments (S+Gh and S+W0) have similarly low expression levels, whereas the two treatments lacking *Spiroplasma* (S0W0 and S0Gh) have similarly high expression levels; implying that there is no influence of *G. hookeri* on expression of these genes. Three of the downregulated genes are involved with male functions (*msl-2, roX1* and *roX2*), one gene (*blanks*) is highly expressed in adult testis, and three other genes (*RpL22-like*, *RpS5b* and *RpS19b*) code for ribosomal proteins (Dataset S4).

At T2, 16 and 106 genes were exclusively up- and down-regulated (respectively) in the S+Gh treatment (exclusive green set in Fig. 4B, Dataset S5). The 16 upregulated genes included three mitochondrially-encoded genes (*ND1, CoIII*, and *Co*I), *necrotic* (which is negative regulator of the Toll pathway; Green et al., 2000), *dumpy* (which is involved in wing development), and one multidrug resistance gene (*Mdr50*). The 106 exclusively downregulated genes did not group in any GO category, making it difficult to link these genes to informative biological functions. The expression values to several of these genes in the S+Gh treatment is similar to S+W0 treatment (Fig S8.). This observation, along with the report that 74 of these genes are highly expressed in adult testis (Dataset S5), suggests that observed expression patterns are likely influenced solely by the presence of *Spiroplasma*. The heatmap of the 16 exclusively upregulated genes (Fig S9), also indicates a possible influence of only *Spiroplasma*. These genes showed a trend of higher expression levels in the S+W0 treatment relative to the S0W0 treatment, but this difference was not significant, possibly due to the high variation among S+W0 replicates.

Only one intersection at T2 (green+blue sets) contained genes (n = 1315; all down-regulated; Fig. 4B). These genes are associated with spermatid differentiation/development functions and 1214 of these 1315 are predominantly expressed in the adult testis (Dataset S5). Thus, their lower expression in the presence of *Spiroplasma* (with or without Gh; see Fig S10) is likely simply the result of an absence of males in these treatments.

### 3.6 Expression of *Spiroplasma* RIP proteins and evidence ribosomal damage

The total number of reads mapped to *Spiroplasma* excluding ribosomal sequences, was very low (range = 1080-17,826 per replicate, Table S2). No DE genes were detected, but this could be a reflection of lack of power. For differential gene expression analyses in bacteria, a minimum of four-five million reads per replicate has been recommended. (Haas et al., 2012) Due to the relevance of RIP genes in *Spiroplasma* (Ballinger and Perlman, 2017), we examined the read counts of the five RIP genes encoded in the *s*Mel genome, which revealed that the RIP2 gene encoding was the most highly expressed at both time points and in all treatments (Figure 5).

**Figure 5.**
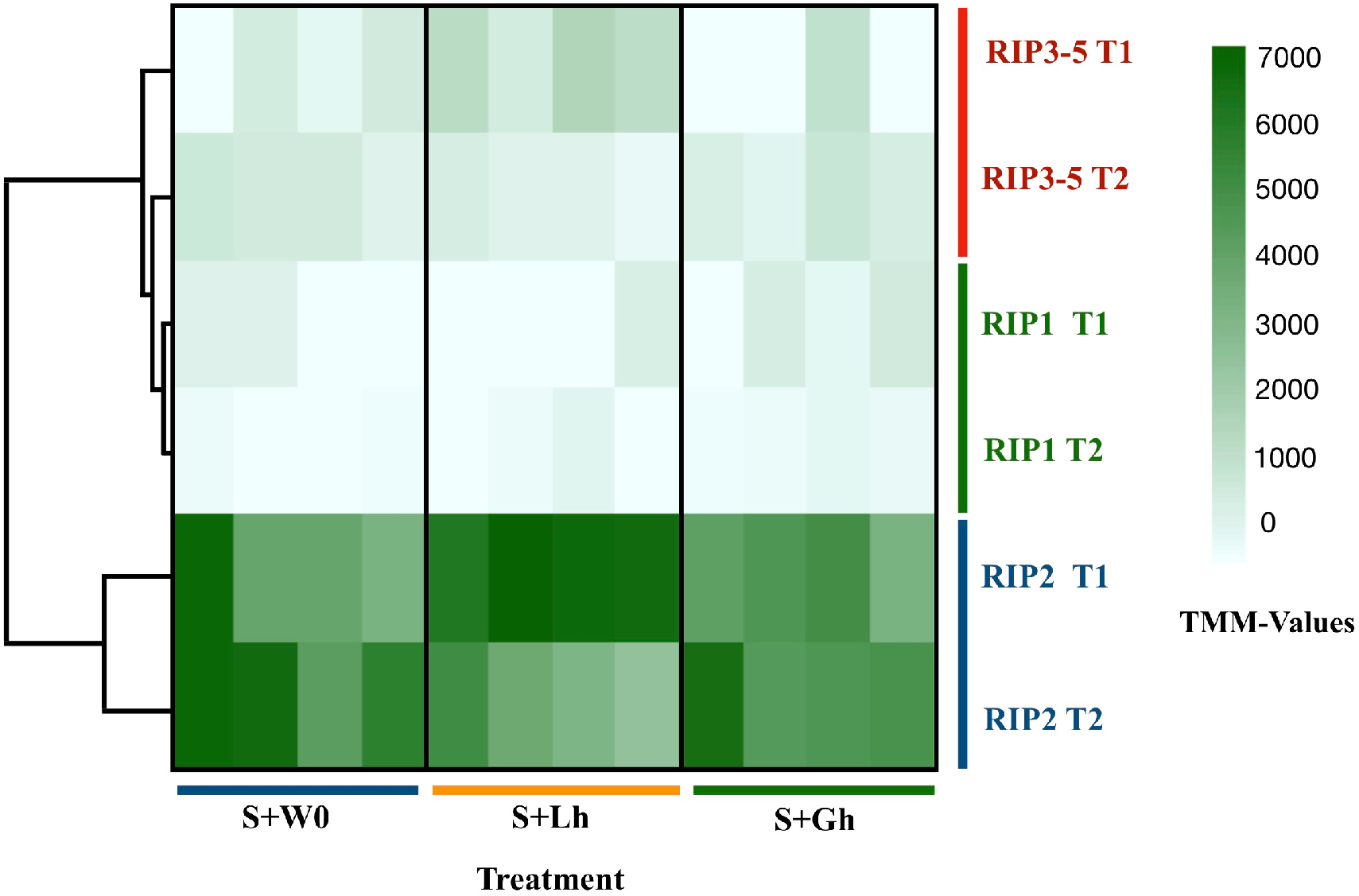
Expression levels of the five ribosome inactivating protein (RIP) genes encoded by Spiroplasma genome, in absence of parasitism (S+W0), in parasitism by *L. heterotoma* (S+Lh) or *G. hookeri* (S0Gh), at 24h (T1) or 72h (T2) post-wasp attack (PWA). Values of TMM edgeR normalized counts are shown. Because the RIP3, RIP4, and RIP5 genes are identical at the nucleotide level, their collective expression levels were reported under the label “RIP3-5”.

To corroborate the observed patterns of RIP gene expression based on the RNA-seq data, RT-qPCR assays were conducted using both the Uganda (UG) and Brazil (BR) *Spiroplasma* sMel strains, and the two wasps, *G. hookeri* and *L. heterotoma*. Consistent with the RNA-seq results, the gene encoding RIP2 was the most highly expressed of the RIP genes in both *Spiroplasma* strains regardless of the presence or absence of wasp (Fig 6, dataset S2). The full output of the statistical analyses is provided in Supplementary file 1. Wasp depurination in the 28S rRNA was significant (X^2^=128.58, df=1, P< 2.2 e-16) in the context of *L. heterotoma* in presence of *Spiroplasma* at T2 (Figure S11 and Table S3). No depurination was detected in *G. hookeri* or fly sequences (Figure S11 and Table S3-4).

**Figure 6.**
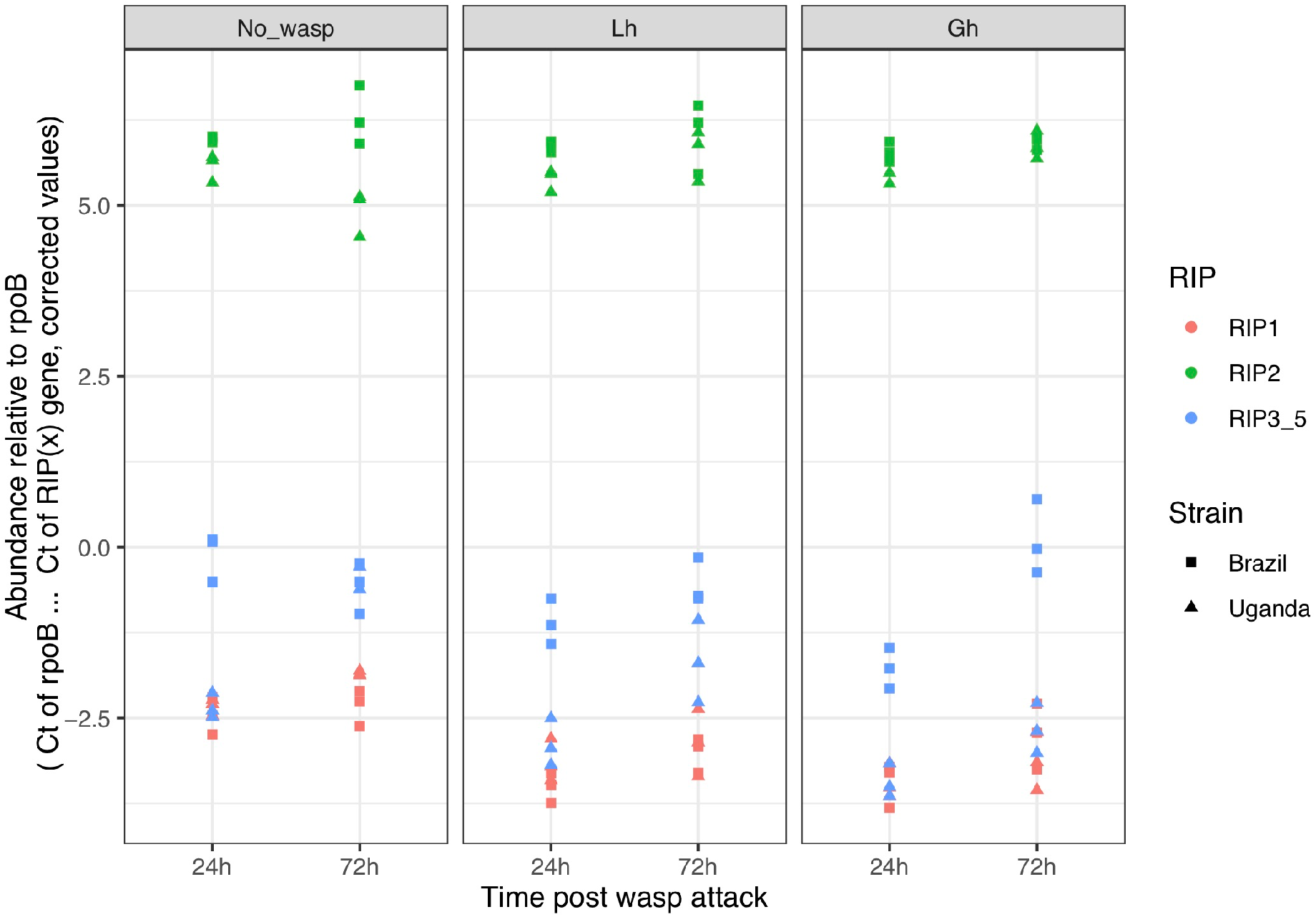
RIP (ribosome inactivating protein) gene expression in *D. melanogaster* larvae in the absence of wasp (No_wasp) or parasitised by *L. heterotoma* (Lh) or *G. hookeri* (Gh) at 24 or 72h post-wasp attack (PWA). Two different S. poulsonii strains were evaluated: sMel-BR and sMel-UG. RIP2 was consistently expressed at a significantly higher level than RIP1 and RIP3-5 in all wasp treatments, time points, and Spiroplasma strains (Tukey HSD P < 0.05).

## 4 Discussion

### 4.1 The adopted ribodepletion procedure is not an ideal strategy for dual transcriptomics in the *Drosophila*-*Spiroplasma* system

In an attempt to recover mRNA reads from both host and symbiont (also known as “dual transcriptome”), we generated RNA-seq libraries avoiding poly-A-tail enrichment step, typically used for eukaryotic mRNA analyses. Instead, our strategy aimed at removal of ribosomal RNA from both host and symbiont by using the RiboZero Epidemiology kit, which is expected to deplete ribosomal RNA from both bacteria and eukaryotes. A previous application of a similar ribodepletion kit, reported removal of > 90% of the rRNA sequences of *D. ananassae* (Kumar et al., 2012). In contrast, in our study the reads that mapped to rRNA genes comprised 3.4–78% of the total reads mapped to *D. melanogaster*, indicating a variable and ineffective degree of ribosomal RNA depletion. Furthermore, by evaluating the pattern of reads mapped to the 28S rRNA gene, we found that ribodepletion effectiveness varied by region of the gene (see Figure S11). We suggest that future applications of ribodepletion in *D. melanogaster* consider including additional probes to capture such regions with higher efficiency. Even if more effective depletion of fly rRNA had been achieved, it is possible that our sequencing effort still would have been inadequate for DE analyses of *Spiroplasma*.

### 4.2 Wasp parasitism in the absence of *Spiroplasma*

The two wasp species used in this study, *L. heterotoma* and *G. hookeri* belong to the same family (Figitidae), but their parasitism strategies are quite different (Schlenke et al., 2007; Mortimer et al., 2013). Our fly transcriptome analysis also revealed differences in the effects of these wasps. Whereas *G. hookeri* appears to activate an immune response in the host, *L. heterotoma* does not, but in turn induces genes related to energy production. Increasing energy could be a strategy of the wasp to obtain more resources from the host. One the other hand, it may reflect energy investment for a fly function such as immune response, which is energetically expensive (Schlenke et al., 2007; Dolezal et al., 2019). If it indeed reflects investment in immune functions, these have not been detected (this study; Schlenke et al., 2007) and are unsuccessful, as the fly generally does not survive attacks by *L. heterotoma* (Xie et al., 2014; Jones and Hurst, 2020b).

The *L. heterotoma*-induced downregulation of genes related to spermatogenesis; a phenomenon not induced by *G. hookeri*, is an interesting contrast between the two wasps. In accordance with our results, (Schlenke et al., 2007) reported downregulation of genes involved with gonad development during parasitism by *L. heterotoma* and *L. boulardi*, albeit at earlier stages. These two independent observations suggest that *L. heterotoma* might induce castration of male hosts. Xie et al., 2011, reported that *Drosophila hydei* males (infected with a non-male-killing strain of *Spiroplasma*) that survived parasitism by *L. heterotoma* (Lh14) had extremely reduced fecundity, but the experiments could not rule out causes other than castration. The phenomenon of male castration has only been reported in parasitoids of the family Braconidae, where the polydnavirus present in the wasp venom induces testis degradation (reviewed in Beckage and Gelman, 2004).

This study is the second one to use a genome-wide approach to evaluate gene expression of *D. melanogaster* in response to *L. heterotoma*. The first one was conducted by Schlenke et al., 2007 and employed microarrays at time points earlier than 24h PWA. In their latest time point (22–24h PWA), Schlenke et al., 2007 found 37 DE genes (*P* < 0.01, −0.5 > FC > 1.0; fold-changes for genes DE at a *P* < 0.05 were not reported). At a similar time point (our T1), we detect only one DE gene (*thor*), which was not detected by Schlenke et al., 2007. Differences between our results and those of Schlenke et al., 2007 may be attributable to the different methodology (e.g. microarrays vs. RNAseq, different software pipelines, and cutoff parameters). Other potential differences could be related to *Drosophila* genotypes or *Wolbachia* infection status, or other experimental conditions or their interactions. In spite of all these variables, both studies provide evidence that parasitism by *L. heterotoma* does not induce an immune response in *D. melanogaster*.

In a different scenario, parasitism by *G. hookeri* induced the Jak/Stat pathway, one of the immune pathways of *D. melanogaster*. Effectors of this pathway (*Tot* and *Tep* genes) were highly upregulated. Tep proteins play an immune role in the mosquito *Anopheles gambiae*, against the parasite *Plasmodium berghei* (Blandin et al., 2004). The function of *Tot* genes is unknown, but they belong to a family of eight genes that are activated by stressful conditions such as bacterial challenge, high temperatures, mechanical pressure, or UV radiation (Ekengren and Hultmark, 2001). Parasitism of *D. melanogaster* by the wasps *L. boulardi* (Schlenke et al., 2007) and *Asobara tabida* (Salazar-Jaramillo et al., 2017) also induce the up-regulation of *Tep* and *Tot* genes, suggesting that upregulation of these genes could be a common response against wasp parasitism. Another potential signal of immune activation against *G. hookeri* is the induction of genes encoding serine proteases, because these enzymes are involved in triggering immune response in insects (reviewed in Alvarado et al., 2020). The melanization cascade, another feature of *D. melanogaster* immune response, also appears to be activated by *G. hookeri*, as two components of this cascade (*PPO3* and *yellow-f*) were up-regulated. Finally the upregulation of genes that are known to be expressed preferentially in hemocytes (i.e., *PPO3*, *ItgaPS4 and he*), or involved in hemocyte proliferation (i.e., *pvf2*) implies that hemocytes are activated by *G. hookeri* parasitism. In accordance with this, Mortimer et al., 2013 showed that the number of lamellocytes increases during parasitism by *G. hookeri*. Even though *D. melanogaster* appears to mount an immune response against *G. hookeri*, the response is inadequate, as ~100% of flies parasitized by this wasp die (Mortimer et al., 2013; Mateos et al., 2016). The reason likely lies in that the venom of this wasp contains a calcium ATPase that prevents activation of plasmatocytes; a necessary early step of the encapsulation process (Mortimer et al., 2013).

### 4.3 Wasp parasitism in the presence of *Spiroplasma*

The primary goal of this study was to identify *Drosophila* fly-encoded functions relevant to the *Spiroplasma*-wasp interaction, as such functions could be involved in the mechanism that leads to the death of certain wasps, including *L. heterotoma*. Our analyses revealed several genes whose expression appears to be influenced specifically by the *Spiroplasma-L. heterotoma* interaction. In the context of the *Spiroplasma*-susceptible wasp (*L. heterotoma*), eight and 32 genes were exclusively DE at T1 and T2, respectively. None of these genes, with exception of attacin-C (*AttC*), were associated with an immune function in *D. melanogaster*. We thus infer that the *Spiroplasma*-mediated wasp-killing mechanism is not strongly influenced by host-encoded immunity, at least at a level detectable by our experiments. Consistent with our findings, an RNA-seq analysis of *Spiroplasma* protection against nematodes in *D. neotestacea* did not detect changes in the host’s immune response (Hamilton et al., 2014). Similarly, Paredes et al., 2016 did not detect an increased cellular response, based on the number of hemocytes, during the *Spiroplasma*-*L. boulardi* interaction. The lack of detectable influence of *Spiroplasma* on host-encoded immunity is consistent with evidence that *Spiroplasma* is not detected as an intruder by the fly, due to the lack of cell wall, where the typical bacterial immune elicitors are found (Hurst et al., 2003; Herren and Lemaitre, 2011).

In the context of *Spiroplasma*-*L. heterotoma* interaction, we identified a set of twelve genes whose expression patterns appear to be “restored”, as well as two genes (TotC and Fbp2) whose expression patterns are likely influenced by the interaction between *Spiroplasma* and *L. heterotoma*. Furthermore, these genes did not exhibit such an expression pattern in the context of the *Spiroplasma*-resistant wasp *G. hookeri*. These genes could contribute to the wasp death and/or fly survival (i.e., a *Spiroplasma*-mediated protection mechanism), but also could simply reflect a side effect of the *Spiroplasma-Lh* interaction. In Group 1 of the restored genes, only two had annotations, one of which is predicted to have chitin-binding activity (*CG42494*) whereas the other has a cytochrome P450 domain (*Cyp4d14*); fold changes of these genes were low (< 2). Cytochrome P450 are a large family of proteins that is involved in detoxification, but also in developmental processes (Chung et al., 2009). For both of these genes, expression levels are reported to go down between the L3 and pupa transition. Therefore, if the S0Lh treatment were developmentally delayed with respect to the S+Lh treatment (see discussion below), our observed expression patterns may simply be a consequence of development time differences between the treatments, rather than a cause of wasp death or enhanced fly survival. Furthermore, if the detoxification function of Cyp4d14 were relevant to the *Spiroplasma*-mediated protection mechanism, we would expect higher expression in the S+Lh than in the S0Lh treatment, which is contrary to our observations. Experimental down-regulation of these and the other Group 1 genes, in the absence of *Spiroplasma*, could help determine whether they have any role in the *Spiroplasma*-mediated protection against wasps.

Genes in Group 2 exhibit a substantial increase in expression in wild type flies during the larva-to-pupa transition according to flybase (Thurmond et al., 2019). Therefore, our observed expression patterns could simply reflect slight differences in fly development time, where *L. heterotoma* slows down host development, but presence of *Spiroplasma* “restores” the development time to that of the unparasitized host. In partial support of this hypothesis, larvae parasitized by *L. hetetoroma* and *L. boulardi* pupate ~2 days later than the unparasitized controls (Schlenke et al., 2007). Whether *Spiroplasma* counteracts the wasp-induced delay in development remains to be determined.

Two genes (*Fbp2* and *TotC*), were not part of the restored genes, but their expression patterns and functions allow us to speculate about their possible roles in the *Spiroplasma-*mediated protection. We acknowledge that the *s*Mel-*L. heterotoma* (strain Lh14) interaction, albeit leading to wasp death, seldom results in fly surviva(Xie et al., 2014; Paredes et al., 2016; Jones and Hurst, 2020b). Given its possible function as a sulfur-storage protein (Meghlaoui and Veuille, 1997), increased expression of *Fbp2* might reflect a greater availability of nutrients for the fly, which might enhance its tolerance to insults from the wasp. Similarly, based on its association to “response to stress”, increased expression of *TotC* may contribute to increasing the fly’s tolerance to the wasp parasitism. In addition, *TotA*, another stress responsive gene, was upregulated but with an FDR value slightly above our cutoff (0.053). Experimental over-expression of these, and the Group 2 “restored” genes, in the absence of *Spiroplasma*, could be used to test whether they influence the outcome of parasitism by *L. heterotoma* and other wasps, including those against which *s*Mel confers stronger rescue (e.g. *L. boulardi* or *L. victoriae*; Mateos et al., 2016; Paredes et al., 2016; Jones and Hurst, 2020a)

### 4.4 RIP expression

Concerning the hypothesis that a *Spiroplasma-*encoded toxin contributes to killing of *Spiroplasma*-susceptible wasps, we found evidence that neither wasp influences expression of RIP genes, but RIP2 gene was highly expressed in contrast to RIP1 or RIP3-5. In a previous report in the absence of parasitism, RIP2 was also found to be the most highly expressed of the RIP genes throughout the fly life cycle (only substrain Uganda was examined; Garcia-Arraez et al., 2019)). The only context where relatively lower expression of RIP2 and RIP1 in *s*Mel has been reported is in the case of in vitro culture versus fly hemolymph (Masson et al., 2018),implying that expression of these genes can be regulated. The expression of RIP genes in context of other tolerant/resistant wasps had not been determined.

Our study also reveals that both the *s*Mel-UG and *s*Mel-BR sub-strains of *S. poulsonii* express their RIP genes at similar levels in the presence or absence of *L. heterotoma* and *G. hookeri*. Although differences in the genomes of substrains *s*Mel-Ug and *s*Mel-BR have been reported (Gerth et al., 2020) our observations suggest that at least for RIP expression patterns the two strains are very similar. Based on expression levels alone, it appears that *s*Mel RIP2 might have a stronger role in wasp death than the other RIP genes. However, it is intriguing that the genome of sHy (the poulsonii-clade native *Spiroplasma* of *D. hydei)* does not harbor a gene with high homology to sMel RIP2. Instead, its genome encodes a gene with high homology to sMel RIP1, as well as putative RIP-encoding genes with homology to genes in *Spiroplasma* strains other than *D. melanogaster* (Gerth et al., 2020), sHy is known to kill *L. heterotoma*, and enhance survival of its host *D. hydei* (Xie et al., 2010). It is thus possible that more than one of the RIP genes in the *Spiroplasma* strains that kill *L. heterotoma*, contributes to wasp killing.

Detection of signals of ribosomal depurination in the *Spiroplasma*-susceptible (*L. heterotoma*) but not in the *Spiroplasma*-resistant (*G. hookeri*) wasp is consistent with the hypothesis that RIP-induced depurination contributes to wasp death. Ballinger and Perlman, 2017, using a more direct approach to evaluate depurination (i.e., qPCR), reported evidence of *Spiroplasma* (strain sMel-UG)-induced depurination in *L. heterotoma*, as well as in *L. boulardi* (another *Spiroplasma*-susceptible wasp), but not in the *Spiroplasma*-resistant pupal ectoparasitic wasp (*Pachycrepoideus vindemmiae*). Ballinger and Perlman, 2017. hypothesized that *P. vindemmiae* resistance to *Spiroplasma* (and depurination) may stem from the fact that it is not immersed in *Spiroplasma*-layden hemolymph during development. This explanation, however, would not apply to *G. hookeri*, which is a larval endo-parasitoid that spends the initial stages of development in the host hemocoel. Whether RIP-induced depurination is necessary and sufficient to kill susceptible wasps has not been determined. Ideally, the effect of RIP on the wasp would be tested in the absence of *Spiroplasma*, or in “knocked-out” mutants in these genes. In addition, the target cells and the mechanism of entry of *Spiroplasma* RIPs has not been determined. It is unclear whether *Spiroplasma* must colonize wasp tissues in order to deliver RIP or if toxins can be acquired during wasp feeding; both RIP1 and RIP2 proteins have been detected in the fly hemolymph (see Garcia-Arraez et al., 2019). It is also possible that other *Spiroplasma*-encoded putative virulence factors such as chitinase (*ChiD*) or glycerol-3-phosphate oxidase (*glpO*) could contribute to wasp death (Masson et al., 2018). Presence of *glpO* in *S. taiwanense* is proposed to be the cause of its pathogenicity to mosquitoes (Lo and Kuo, 2017).

Assuming RIP is an important factor in wasp killing, the apparent lack of ribosome depurination of G. hookeri, along with the unaltered expression of RIP genes, suggest that this wasp avoids RIP-induced damage by interfering with translation of RIP mRNA, or by inactivating RIP. Resistance to RIP toxicity has been reported in Lepidoptera, and has been attributed to serine protease-mediated hydrolysis in the digestive tract (Gatehouse et al., 1990). As suggested by Ballinger and Perlman, 2017 for *P. vindemmiae*, it is possible that conditions in the gut of *G. hookeri* inactivate ingested RIPs. Alternatively, RIP proteins may be active but unable to enter *G. hookeri* cells, or unable to reach the appropriate cellular compartments to damage ribosomes. It is unlikely that wasp ribosomes that come into contact with RIPs are immune to depurination because of the extremely conserved eukaryotic motif targeted by these proteins.

## Conclusions

We detected a small number of fly genes whose expression was influenced by the interaction between *Spiroplasma* and the susceptible wasp *L. heterotoma*. Based on the annotations of these genes, our results suggest that immune priming does not contribute to the *Spiroplasma*-mediated wasp killing mechanism. Furthermore, based on the reported expression patterns of several of these genes during fly development, it is possible that our results reflect *Spiroplasma*-mediated changes in development time of Lh-parasitized flies, which has yet to be verified. In accordance with previous studies, we detected evidence consistent with ribosomal damage in the susceptible but not the resistant wasp. In addition, our transcriptome and qPCR results indicated that RIP2 is the most highly expressed of the five *s*Mel-encoded RIP genes in both the BR and UG substrains, and that its expression levels were independent of wasp parasitism. We recommend that future studies examine whether any of the sMel-encoded RIPs are necessary and sufficient for wasp killing. The observation that the *Spiroplasma*-resistant wasp *G. hookeri* did not influence RIP transcript levels raises interesting questions about its mechanism of resistance or tolerance to *Spiroplasma*.

## Acknowledgements

Texas Agrilife Genomics and Bioinformatics Services for library preparation, sequencing, and advice on handling of RNA. Luis E Servín-Garcidueñas critically reviewed the manuscript. Renato D. La Torre Ramirez for help with qPCR standard curve. Alfredo Hernández, Víctor del Moral and Romualdo Zayas for the provided computing support.

All bioinformatic analyses were performed on CCG-UNAM servers. Victor Manuel Higareda Alvear is a doctoral student at Programa de Doctorado en Ciencias Biomédicas of the Universidad Nacional Autónoma de México (UNAM), and was supported by Consejo Nacional de Ciencia y Tecnología (CONACyT, CVU 446829).

## References

Agaisse, H., and Perrimon, N. (2004). The roles of JAK/STAT signaling in Drosophila immune responses. Immunol. Rev. 198, 72–82. doi:10.1111/j.0105-2896.2004.0133.x.

Alvarado, G., Holland, S. R., DePerez-Rasmussen, J., Jarvis, B. A., Telander, T., Wagner, N., et al. (2020). Bioinformatic analysis suggests potential mechanisms underlying parasitoid venom evolution and function. Genomics 112, 1096–1104. doi:10.1016/j.ygeno.2019.06.022.

Andrews, S. (2010). FastQC: A Quality Control Tool for High Throughput Sequence Data [Online]. Available online at: http://www.bioinformatics.babraham.ac.uk/projects/fastqc/

Ballinger, M. J., and Perlman, S. J. (2017). Generality of toxins in defensive symbiosis: Ribosome-inactivating proteins and defense against parasitic wasps in Drosophila. PLoS Pathog. 13, 1–19. doi:10.1371/journal.ppat.1006431.

Beckage, N. E., and Gelman, D. B. (2004). Wasp Parasitoid Disruption of Host Development: Implications for New Biologically Based Strategies for Insect Control. Annu. Rev. Entomol. 49, 299–330. doi:10.1146/annurev.ento.49.061802.123324.

Blandin, S., Shiao, S. H., Moita, L. F., Janse, C. J., Waters, A. P., Kafatos, F. C., et al. (2004). Complement-like protein TEP1 is a determinant of vectorial capacity in the malaria vector Anopheles gambiae. Cell 116, 661–670. doi:10.1016/S0092-8674(04)00173-4.

Bolger, A. M., Lohse, M., and Usadel, B. (2014). Trimmomatic: A flexible trimmer for Illumina sequence data. Bioinformatics 30, 2114–2120. doi:10.1093/bioinformatics/btu170.

Brandt, J. W., Chevignon, G., Oliver, K. M., and Strand, M. R. (2017). Culture of an aphid heritable symbiont demonstrates its direct role in defence against parasitoids. Proc. R. Soc. B Biol. Sci. 284. doi:10.1098/rspb.2017.1925.

Bray, N. L., Pimentel, H., Melsted, P., and Pachter, L. (2016). Near-optimal probabilistic RNA-seq quantification. Nat. Biotechnol. 34, 525–527. doi:10.1038/nbt.3519.

Caragata, E. P., Rancés, E., Hedges, L. M., Gofton, A. W., Johnson, K. N., O’Neill, S. L., et al. (2013). Dietary Cholesterol Modulates Pathogen Blocking by Wolbachia. PLoS Pathog. 9. doi:10.1371/journal.ppat.1003459.

Carton, Y., Poirié, M., and Nappi, A. J. (2008). Insect immune resistance to parasitoids. Insect Sci. 15, 67–87. doi:10.1111/j.1744-7917.2008.00188.x.

Cheng, B., Kuppanda, N., Aldrich, J. C., Akbari, O. S., and Ferree, P. M. (2016). Male-killing Spiroplasma alters behavior of the dosage compensation complex during drosophila melanogaster embryogenesis. Curr. Biol. 26, 1339–1345. doi:10.1016/j.cub.2016.03.050.

Chung, H., Sztal, T., Pasricha, S., Sridhar, M., Batterham, P., and Daborn, P. J. (2009). Characterization of Drosophila melanogaster cytochrome P450 genes. Proc. Natl. Acad. Sci. U. S. A. 106, 5731–5736. doi:10.1073/pnas.0812141106.

Corbin, C., Jones, J. E., Chrostek, E., Fenton, A., and Hurst, G. D. D. (2020). Thermal sensitivity of the Spiroplasma-Drosophila hydei protective symbiosis: The best of climes, the worst of climes. bioRxiv. doi:10.1101/2020.04.30.070938.

Dolezal, T., Krejcova, G., Bajgar, A., Nedbalova, P., and Strasser, P. (2019). Molecular regulations of metabolism during immune response in insects. Insect Biochem. Mol. Biol. 109, 31–42. doi:10.1016/j.ibmb.2019.04.005.

Dudzic, J. P., Kondo, S., Ueda, R., Bergman, C. M., and Lemaitre, B. (2015). Drosophila innate immunity: regional and functional specialization of prophenoloxidases. BMC Biol. 13, 81. doi:10.1186/s12915-015-0193-6.

Ekengren, S., and Hultmark, D. (2001). A family of Turandot-related genes in the humoral stress response of Drosophila. Biochem. Biophys. Res. Commun. 284, 998–1003. doi:10.1006/bbrc.2001.5067.

Garcia-Arraez, M. G., Masson, F., Escobar, J. C. P., and Lemaitre, B. (2019). Functional analysis of RIP toxins from the Drosophila endosymbiont Spiroplasma poulsonii. BMC Microbiol. 19, 1–10. doi:10.1186/s12866-019-1410-1.

Gatehouse, A. M. R., Barbieri, L., Stirpe, F., and Croy, R. R. D. (1990). Effects of ribosome inactivating proteins on insect development – differences between Lepidoptera and Coleoptera. Entomol. Exp. Appl. 54, 43–51. doi:10.1111/j.1570-7458.1990.tb01310.x.

Gerth, M., Humberto., M.-M., Ramirez, P., Masson, F., Griffin, J., Aramayo, R., et al. (2020). Rapid molecular evolution of Spiroplasma symbionts of Drosophila. bioRxiv. doi:10.1101/2020.06.23.165548.

Green, C., Levashina, E., McKimmie, C., Dafforn, T., Reichhart, J. M., and Gubb, D. (2000). The necrotic gene in Drosophila corresponds to one of a cluster of three serpin transcripts mapping at 43A1.2. Genetics 156, 1117–1127.

Haas, B. J., Chin, M., Nusbaum, C., Birren, B. W., and Livny, J. (2012). How deep is deep enough for RNA-Seq profiling of bacterial transcriptomes? BMC Genomics 13, 734. doi:10.1186/1471-2164-13-734.

Hamilton, P. T., Leong, J. S., Koop, B. F., and Perlman, S. J. (2014). Transcriptional responses in a Drosophila defensive symbiosis. Mol. Ecol. 23, 1558–1570. doi:10.1111/mec.12603.

Hamilton, P. T., Peng, F., Boulanger, M. J., and Perlman, S. J. (2015). A ribosome-inactivating protein in a Drosophila defensive symbiont. Proc Natl Acad Sci U S A, 1518648113-. doi:10.1073/pnas.1518648113.

Han, Q., Fang, J., Ding, H., Johnson, J. K., Christensen, B. M., and Li, J. (2002). Identification of Drosophila melanogaster yellow-f and yellow-f2 proteins as dopachrome-conversion enzymes. Biochem. J. 368, 333–340. doi:10.1042/BJ20020272.

Hart, S. N., Therneau, T. M., Zhang, Y., Poland, G. A., and Kocher, J. P. (2013). Calculating sample size estimates for RNA sequencing data. J. Comput. Biol. 20, 970–978. doi:10.1089/cmb.2012.0283.

Harumoto, T., Anbutsu, H., Lemaitre, B., and Fukatsu, T. (2016). Male-killing symbiont damages host’s dosage-compensated sex chromosome to induce embryonic apoptosis. Nat. Commun. 7, 12781. doi:10.1038/ncomms12781.

Herren, J. K., and Lemaitre, B. (2011). Spiroplasma and host immunity: Activation of humoral immune responses increases endosymbiont load and susceptibility to certain Gram-negative bacterial pathogens in Drosophila melanogaster. Cell. Microbiol. 13, 1385–1396. doi:10.1111/j.1462-5822.2011.01627.x.

Hillyer, J. F. (2016). Insect immunology and hematopoiesis. Dev. Comp. Immunol. 58, 102–118. doi:10.1016/j.dci.2015.12.006.

Hurst, G. D. D., Anbutsu, H., Kutsukake, M., and Fukatsu, T. (2003). Hidden from the host: Spiroplasma bacteria infecting Drosophila do not cause an immune response, but are suppressed by ectopic immune activation. Insect Mol. Biol. 12, 93–97. doi:10.1046/j.1365-2583.2003.00380.x.

Irving, P., Ubeda, J. M., Doucet, D., Troxler, L., Lagueux, M., Zachary, D., et al. (2005). New insights into Drosophila larval haemocyte functions through genome-wide analysis. Cell. Microbiol. 7, 335–350. doi:10.1111/j.1462-5822.2004.00462.x.

Jaenike, J., Unckless, R., Cockburn, S. N., Boelio, L. M., and Perlman, S. J. (2010). Adaptation via symbiosis: Recent spread of a drosophila defensive symbiont. Science (80-.). 329, 212–215. doi:10.1126/science.1188235.

Jones, J. E., and Hurst, G. D. D. (2020a). Symbiont-mediated fly survival is independent of defensive symbiont genotype in the Drosophila melanogaster-Spiroplasma-wasp interaction. bioRxiv, 1–24. doi:10.1101/2020.07.05.154906.

Jones, J. E., and Hurst, G. D. D. (2020b). Symbiont-mediated protection varies with wasp genotype in the Drosophila melanogaster–Spiroplasma interaction. Heredity (Edinb). 124, 592–602. doi:10.1038/s41437-019-0291-2.

Kim, D., Langmead, B., and Salzberg, S. L. (2015). HISAT: A fast spliced aligner with low memory requirements. Nat. Methods 12, 357–360. doi:10.1038/nmeth.3317.

Kumar, N., Creasy, T., Sun, Y., Flowers, M., Tallon, L. J., and Dunning Hotopp, J. C. (2012). Efficient subtraction of insect rRNA prior to transcriptome analysis of Wolbachia-Drosophila lateral gene transfer. BMC Res. Notes 5, 1. doi:10.1186/1756-0500-5-230.

Liao, Y., Smyth, G. K., and Shi, W. (2014). FeatureCounts: An efficient general purpose program for assigning sequence reads to genomic features. Bioinformatics 30, 923–930. doi:10.1093/bioinformatics/btt656.

Lo, W. S., and Kuo, C. H. (2017). Horizontal Acquisition and Transcriptional Integration of Novel Genes in Mosquito-Associated Spiroplasma. Genome Biol. Evol. 9, 3246–3259. doi:10.1093/gbe/evx244.

Lucchesi, J. C., and Kuroda, M. I. (2015). Dosage compensation in drosophila. Cold Spring Harb. Perspect. Biol. 7, 1–21. doi:10.1101/cshperspect.a019398.

Lyne, R., Smith, R., Rutherford, K., Wakeling, M., Varley, A., Guillier, F., et al. (2007). FlyMine: An integrated database for Drosophila and Anopheles genomics. Genome Biol. 8. doi:10.1186/gb-2007-8-7-r129.

Masson, F., Copete, S. C., Schüpfer, F., Garcia-Arraez, G., and Lemaitre, B. (2018). In Vitro culture of the insect Endosymbiont Spiroplasma poulsonii highlights bacterial genes involved in host-symbiont interaction. MBio 9, 1–11. doi:10.1128/mBio.00024-18.

Mateos, M., Winter, L., Winter, C., Higareda-Alvear, V. M., Martinez-Romero, E., and Xie, J. (2016). Independent origins of resistance or susceptibility of parasitic wasps to a defensive symbiont. Ecol. Evol., 2679–2687. doi:10.1002/ece3.2085.

Meghlaoui, G. K., and Veuille, M. (1997). Selection and methionine accumulation in the fat body protein 2 gene (FBP2), a duplicate of the Drosophila alcohol dehydrogenase (ADH) gene. J. Mol. Evol. 44, 23–32. doi:10.1007/PL00006118.

Montenegro, H., Solferini, V. N., Klaczko, L. B., and Hurst, G. D. D. (2005). Male-killing Spiroplasma naturally infecting Drosophila melanogaster. Insect Mol. Biol. 14, 281–287. doi:10.1111/j.1365-2583.2005.00558.x.

Mortimer, N. T., Goecks, J., Kacsoh, B. Z., Mobley, J. A., and Bowersock, G. J. (2013). Parasitoid wasp venom SERCA regulates Drosophila calcium levels and inhibits cellular immunity. 110, 9427–9432. doi:10.1073/pnas.1222351110.

Oliver, K. M., and Perlman, S. J. (2020). Toxin-mediated protection against natural enemies by insect defensive symbionts. 1st ed., eds. K. Oliver and Russel J. Elsevier Ltd. doi:10.1016/bs.aiip.2020.03.005.

Oliver, K. M., Russell, J. A., Morant, N. A., and Hunter, M. S. (2003). Facultative bacterial symbionts in aphids confer resistance to parasitic wasps. Proc. Natl. Acad. Sci. U. S. A. 100, 1803–1807. doi:10.1073/pnas.0335320100.

Paredes, J. C., Herren, J. K., Schüpfer, F., and Lemaitre, B. (2016). The Role of Lipid Competition for Endosymbiont-Mediated Protection against Parasitoid Wasps in *Drosophila*. MBio 7, e01006–16. doi:10.1128/mBio.01006-16.

Pool, J. E., Wong, A., and Aquadro, C. F. (2006). Finding of male-killing Spiroplasma infecting Drosophila melanogaster in Africa implies transatlantic migration of this endosymbiont. Heredity (Edinb). 97, 27–32. doi:10.1038/sj.hdy.6800830.

Riegler, M., Sidhu, M., Miller, W. J., and O’Neill, S. L. (2005). Evidence for a global Wolbachia replacement in Drosophila melanogaster. Curr. Biol. 15, 1428–1433. doi:10.1016/j.cub.2005.06.069.

Robinson, M. D., McCarthy, D. J., and Smyth, G. K. (2009). edgeR: A Bioconductor package for differential expression analysis of digital gene expression data. Bioinformatics 26, 139–140. doi:10.1093/bioinformatics/btp616.

Russo, J. (1996). Insect immunity: Early events in the encapsulation process of parasitoid (Leptopilina boulardi) eggs in resistant and susceptible strains of Drosophila. Parasitology 112, 135–142. doi:10.1017/s0031182000065173.

Salazar-Jaramillo, L., Jalvingh, K. M., de Haan, A., Kraaijeveld, K., Buermans, H., and Wertheim, B. (2017). Inter- and intra-species variation in genome-wide gene expression of Drosophila in response to parasitoid wasp attack. BMC Genomics 18, 1–14. doi:10.1186/s12864-017-3697-3.

Schlenke, T. A., Morales, J., Govind, S., and Clark, A. G. (2007). Contrasting infection strategies in generalist and specialist wasp parasitoids of Drosophila melanogaster. PLoS Pathog. 3, 1486–1501. doi:10.1371/journal.ppat.0030158.

Stirpe, F. (2004). Ribosome-inactivating proteins. Toxicon 44, 371–383. doi:10.1016/j.toxicon.2004.05.004.

Thurmond, J., Goodman, J. L., Strelets, V. B., Attrill, H., Gramates, L. S., Marygold, S. J., et al. (2019). FlyBase 2.0: The next generation. Nucleic Acids Res. 47, D759–D765. doi:10.1093/nar/gky1003.

Wang, J., Wu, Y., Yang, G., and Aksoy, S. (2009). Interactions between mutualist Wigglesworthia and tsetse peptidoglycan recognition protein (PGRP-LB) influence trypanosome transmission. Proc. Natl. Acad. Sci. U. S. A. 106, 12133–12138. doi:10.1073/pnas.0901226106.

Wey, B., Heavner, M. E., Wittmeyer, K. T., Briese, T., Hopper, K. R., and Govind, S. (2020). Immune suppressive extracellular vesicle proteins of Leptopilina heterotoma are encoded in the wasp genome. G3 Genes, Genomes, Genet. 10, 1–12. doi:10.1534/g3.119.400349.

Xie, J., Butler, S., Sanchez, G., and Mateos, M. (2014). Male killing Spiroplasma protects Drosophila melanogaster against two parasitoid wasps. Heredity (Edinb). 112, 399–408. doi:10.1038/hdy.2013.118.

Xie, J., Tiner, B., Vilchez, I., and Mateos, M. (2011). Effect of the Drosophila endosymbiont Spiroplasma on parasitoid wasp development and on the reproductive fitness of wasp-attacked fly survivors. Evol. Ecol. 25, 1065–1079. doi:10.1007/s10682-010-9453-7.

Xie, J., Vilchez, I., and Mateos, M. (2010). Spiroplasma bacteria enhance survival of Drosophila hydei attacked by the parasitic wasp Leptopilina heterotoma. PLoS One 5. doi:10.1371/journal.pone.0012149.

